# Shavenbaby and Yorkie mediate Hippo signaling to protect adult stem cells from apoptosis

**DOI:** 10.1101/163279

**Authors:** Jérôme Bohère, Alexandra Mancheno-Ferris, Kohsuke Akino, Yuya Yamabe, Sachi Inagaki, Hélène Chanut-Delalande, Serge Plaza, Yuji Kageyama, Dani Osman, Cédric Polesello, François Payre

## Abstract

To compensate for accumulating damages and cell death, adult homeostasis (e.g., body fluids and secretion) requires organ regeneration, operated by long-lived stem cells. How stem cells can survive throughout the animal life yet remains poorly understood. Here we show that the transcription factor Shavenbaby (Svb, OvoL in vertebrates) is expressed in renal/nephric stem cells (RNSCs) of *Drosophila* and required for their maintenance during adulthood. As recently shown in embryos, Svb function in adult RNSCs further needs a post-translational processing mediated by Polished rice (Pri) smORF peptides and impairing Svb function leads to RNSC apoptosis. We show that Svb interacts both genetically and physically with Yorkie (YAP/TAZ in vertebrates), a nuclear effector of the Hippo pathway, to activate the expression of the inhibitor of apoptosis *DIAP1*. These data therefore identify Svb as a novel nuclear effector in the Hippo pathway, critical for the survival of adult somatic stem cells.

---

The family of OvoL/Ovo/Shavenbaby (Svb) transcription factors has been strongly conserved across evolution^1^ and is characteristic of animal species. Initially discovered in flies for a dual function in the development of epidermal derivatives (Svb) and of the germline (Ovo)^2, 3^, mammalian orthologs (OvoL1-3) have soon been identified^4–6^. *OvoL*/*svb* genes produce several protein isoforms and the existence of three partially redundant paralogs in mammals complicates their genetic analysis. There is a single gene in *Drosophila*, which expresses germline- (*ovo*) and somatic-specific (*svb*) transcripts from different promoters. Previous work has well-established the role of Svb in the development of embryonic epidermal tissues^3^, where it triggers a tridimensional cell shape remodeling for the formation of actin-rich apical extensions, called trichomes. *Svb* expression is driven by a large array of *cis*-regulatory regions, which have become a fruitful paradigm for elucidating the function^7, 8^ and evolution^9–11^ of developmental enhancers. Svb enhancers directly integrate multiple inputs form upstream regulatory pathways^7^ and often drive similar patterns, all together conferring robustness to epidermal development in the face of varying environmental conditions and/or genetic backgrounds^7, 8^. During embryogenesis, the Svb transcription factor directly activates a battery of >150 target genes^12–14^ collectively responsible for actin and extra-cellular-matrix reorganization that underlie trichome formation^15^. Recent studies have unraveled a tight control of Svb transcriptional properties, in response to Polished rice (Pri, also known as Tarsal-less) peptides, which belongs to a novel family of peptides encoded from small open reading frames (smORF) hidden within apparently long noncoding RNAs^16^. Svb is first translated as a long-sized protein that acts as a repressor (Svb^REP^)^17^. Pri smORF peptides then induce a proteolytic processing of Svb^REP^ leading to the degradation of its N-terminal region and releasing a shorter activator form, Svb^ACT 17^ Further work has demonstrated that *pri* expression is directly regulated by periodic pulses of steroid hormones^18^, allowing a functional connection between hard-wired genetic regulatory networks (*svb* expression) and systemic hormonal control (mediated by *pri*) for a proper spatio-temporal control of epidermal cell morphogenesis^16^.

Recent studies suggest that OvoL/Svb factors display broader functions throughout epithelial tissues in both normal and various pathological situations. Molecular profiling of human tumors has revealed that OvoL deregulation is a feature of many carcinomas, directly linked to the metastatic potential of morbid cancers^19–23^, including kidney^24^. OvoL factors have been proposed^25, 26^ to counteract a conserved core of regulators composed of Snail/Slug and Zeb1-2 transcription factors, as well as the micro RNA *mir200,* well known to promote epithelial-mesenchymal transition (EMT)^27^. The activity of OvoL might help stabilizing a hybrid E/M phenotype^21, 25^, providing many advantages for both tumors and normal stem cells^28^. Indeed, recent data show that, like adult somatic stem cells, the most aggressive tumors often display a hybrid phenotype between mesenchymal and epithelial states^27^, and the expression of specific OvoL isoforms can annihilate the metastatic potential of mammary tumors^29,19^. In addition, OvoL/Svb factors have been linked to the control of various progenitors/stem cells, from basal invertebrates^30^ to mammals^31–33^. Therefore, whereas a large body of evidence supports a key role for OvoL/Svb in the behavior of somatic stem cells, a functional investigation of their mode of action *in vivo* remains to be undertaken.

Here we built on the knowledge and tools accumulated for the study of Svb function in flies to investigate its putative contribution to the behavior of somatic stem cells in the adult. We show that in the Malpighian tubules, which ensure essential renal functions in insects^34, 35^, *svb* is specifically expressed in the adult renal and nephric stem cells (RNSCs, see Fig. 1a). We further find that a main function of Svb in the kidney is to protect RNSCs from apoptosis by controlling the expression of the inhibitor of apoptosis, *DIAP1,* in interaction with Yorkie, a nuclear effector of the Hippo pathway.

**Figure 1:**
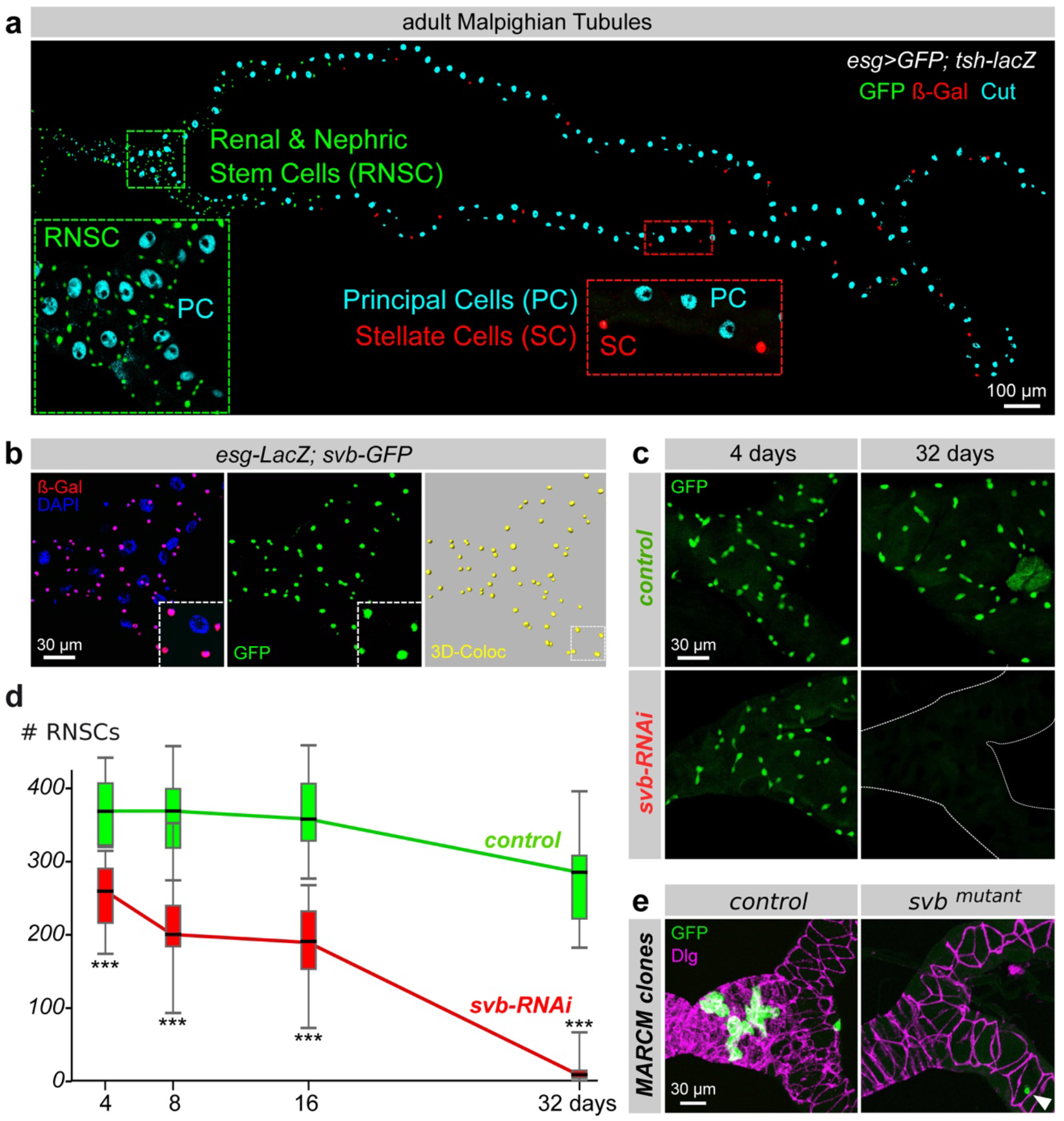
*svb* is expressed in RNSCs and required for their maintenance. **(a)** Adult Malpighian tubules are composed of three types of cells. Principal cells (PC) are identified by immunostaining against Cut (cyan) and stellate cells (SC) by *tsh-LacZ* (in red). RNSCs located in the lower tubules express *esg-Gal4*, *UAS-GFP* (green). **(b)** Fork region of the Malpighian tubules. The expression of *svb*and *esg* was monitored by the expression of *svb-E-GFP* (green) and *esg-LacZ* (red) enhancers, respectively. Nuclei were stained with DAPI (blue). **(c)** *esg*^*ts*^-driven *svb-RNAi* leads to a progressive decrease in the number of RNSCs compared to control conditions (*esg*^*ts*^ driving only GFP). **(d)** Quantification of the number of RNSCs (*esg-GFP* positive) after 4, 8, 16 and 32 of transgene induction in control (green) and *svb-RNAi* (red) conditions. **(e)** Genetic mosaics (MARCM) showing control and *svb*^*R9*^ clones, positively labelled with GFP (green) in the fork region of Malpighian tubules, 25 days after clone induction. The white arrowhead indicates the position of a svb-mutant cell.

## Results and Discussion

### *svb* is expressed in Renal Nephric Stem Cells and controls their maintenance

To assay whether *svb* might be expressed in the adult, we tested large genomic reporter constructs that cover each of the seven enhancers contributing to *svb* expression. We found that one enhancer^9^, *svb*^*E10*^, drove specific expression in tiny cells of the Malpighian tubules (Supplementary Fig. 1a, b).

Adult Malpighian tubules are mainly composed of two types of differentiated cells^35^ (Fig. 1a). The principal cells -characterized by the homeodomain Cut protein- express the vacuolar-ATPase (V-ATPase) that establishes an H^+^ electrochemical potential promoting *trans-epithelial* secretion of Na^+^ and K^+^ ^34^. The second main population of Malpighian tubules are termed stellate cells, featured by the expression of the Teashirt transcription factor, and that regulate the transport of Cl^−^ and water^34^. While both principal and stellate cells display large-sized polyploid nuclei, a third population of tiny diploid cells^36^ are located in the lower tubules and correspond to RNSCs^34, 35, 37^ (Fig. 1a). RNSCs derive from a subpopulation of intestinal stem cell precursors that colonize Malpighian tubules during post-embryonic development^37, 38^. RNSCs are characterized by the expression of Escargot, a transcription factor of the Snail/SLUG family that is also expressed in intestinal stem cells (ISCs^39^) where it acts to prevent ISC differentiation^40, 41^. Co-localization with an *esg-LacZ* reporter confirmed that the *svb*^*E10*^ enhancer was active in RNSCs (Fig. 1b and Supplementary Fig. 1b). To define the minimal region of *svb* responsible for the expression in RNSCs, we assayed a collection of overlapping constructs^7^. This identified two independent elements, the *svb*^*E3N*^ and *svb*^*E6*^ enhancers^7, 9^, which despite having distinct activities during embryogenesis^9, 42^ drove similar expression in adult RNSCs (Supplementary Fig. 1c).

Having established that two enhancers drive specific expression of *svb* in the adult stem cells of the renal system, we next assayed consequences of depleting *svb* function in RNSCs. We used a well-controlled genetic system, hereafter referred to as *esg*^*ts*^, ensuring RNAi-mediated gene depletion, specifically in the stem cells and only at the adult stage^43^. We also developed an image analysis pipeline, allowing automated quantification of the whole population of RNSCs (see methods). In control conditions, the number of *esg*-positive RNSCs remains stable after adult hatching, with approx. 350 cells *per* tubules (Fig. 1c,d). We only noticed a weak reduction of RNSCs (300 cells) after one month. In contrast, *esg*^*ts*^-driven RNAi depletion of *svb* led to a dramatic loss of RNSCs, which were completely absent after 32 days of treatment (Fig. 1c,d). The effects of *svb* depletion were already strong following 8 days of treatment, with a two-fold reduction in the number of RNSCs. Similar results were observed when using an independent driver of RNSCs (*Dome-meso-gal4)* to direct RNAi-mediated knockdown of *svb* (Supplementary Fig. 1d,e). The loss of RNSCs upon *svb* depletion was also confirmed by staining against Hindsight, another transcription factor specific of RNSCs within Malpighian tubules (Supplementary Fig. 1d,e). Finally, the key role of *svb* in the maintenance of adult RNSCs was further demonstrated by results from genetic mosaics, showing that *svb*-null mutant cells^44^ were unable to maintain RNSCs (Fig. 1e).

Taken together, these data thus reveal that *svb* is specifically expressed in RNSCs and critically required for the maintenance of the adult stem cell compartment.

### Svb processing is essential for its activity in Renal Nephric Stem Cells

In the epidermis, Svb activity relies on a proteolytic processing that causes a switch from a repressor to an activator form. This processing is gated by Pri regulatory peptides, which bind to and activate the Ubr3 ubiquitin E3-ligase that, in turn, triggers a limited degradation operated by the proteasome^45^ (Fig. 2a). Thereby, *pri* mediates a systemic control of Svb maturation since the expression of *pri* is directly regulated by the ecdysone receptor (EcR)^18^.

**Figure 2:**
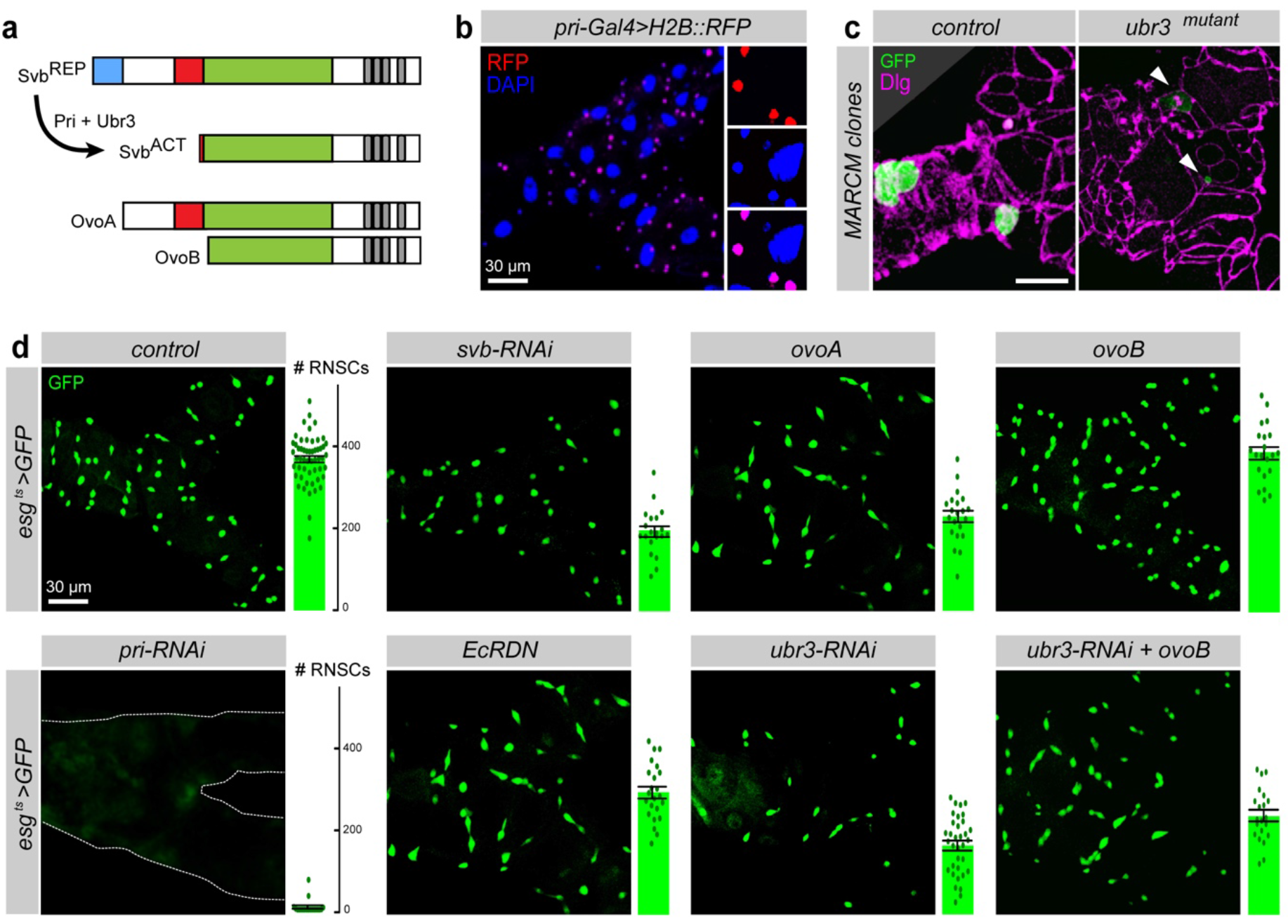
Processing of Svb is essential for RNSC maintenance. **(a)** Schematic representation of Svb maturation, as well as the germinal isoforms OvoA and OvoB that act as constitutive (*pri-* independent) repressor and activator, respectively. **(b)** Expression of *pri* monitored by the activity of *pri- Gal4* driving the expression of H2B::RFP (red). The nuclei (DAPI) are in blue. **(c)** MARCM clones of control and *ubr3*^*B*^ cells, 25 days after induction. Arrowheads indicate positions of *ubr3* mutant cells. **(d)** Fork region of Malpighian tubules, with *esg*^*ts*^-driven expression of GFP and the indicated transgenes, after 8 days of induction. Quantification of the number of *esg+* cells *per* tubule is indicated on the right of each picture (see also Supplementary Figure 2).

To assess whether the function of Svb in Malpighian tubules also required its proteolytic maturation, we investigated a putative function of *pri* and *ubr3* in RNSCs. We screened a collection of *pri* reporter lines^18, 46^ and identified two *cis*-regulatory regions driving expression in RNSCs (Fig. 2b and Supplementary Fig. 2a,b). Consistently with the expression of *pri* in RNSCs, *pri* depletion almost fully eliminated RNSCs upon 8 days of RNAi treatment (Fig. 2d and Supplementary Fig. 2c). In addition, a dominant negative form of the Ecdysone Receptor (EcRDN) that abolishes *pri* expression during both embryonic and post- embryonic development^18^ was sufficient to reduce the number of stem cells when specifically expressed in adult RNSCs (Fig. 2d). Furthermore, we found that *ubr3* was also required for RNSC maintenance, as deduced from results of RNAi-mediated depletion and genetic nullification^45^ of *ubr3* activity (Fig. 2c,d). Finally, the expression of OvoA that behaves as a constitutive repressor isoform of Svb^17, 44, 47^ mimicked the effects observed in *svb* loss of function conditions (Fig. 2d). Reciprocally, the expression of OvoB that acts as a constitutive activator isoform of Svb^17, 44, 47^ was sufficient to rescue the lack of *ubr3* function (Fig. 2d and Supplementary Fig. 2c), demonstrating that Svb function in RNSCs relies on its matured transcription activator form.

These results provide compelling evidence that the whole regulatory machinery discovered for its role in the development of epidermal cells^17, 18, 45^ is also at work in adult RNSCs. We therefore concluded that the post-translational maturation of the Svb transcription factor is essential for the maintenance of RNSCs.

### Svb protects Renal Nephric Stem Cells from apoptosis

The loss of RNSCs observed following the lack of *svb* function or maturation could theoretically result from at least three different causes: i) lack of proliferation, ii) precocious differentiation, or iii) increased cell death. Consistent with the quiescent behavior of RNSCs, we observed a very low frequency of RNSC mitosis in controls, as deduced from staining with the mitotic marker Histone3-P (Supplementary Fig. 3a) and as previously noticed^37^. Therefore, even a complete block of stem cell division cannot account for the disappearance of RNSCs observed in the absence of *svb.* Using the lineage-tracing system called ReDDM that has been recently developed for intestinal stem cells^48^, we next investigated a putative influence of *svb* on RNSC differentiation. Based on differences in the stability of two fluorescent proteins, ReDDM allows marking renal progenitors that express both mCD8::GFP and H2B::RFP, while their progeny only maintain the very stable H2B::RFP ^48^. In control conditions, we detected very rare H2B::RFP progeny (Fig. 3a) confirming low cell renewal in Malpighian tubules^37^. Recent work has shown that the expression of *mir-8* (the *Drosophila* homolog of *mir-200* in vertebrates) downregulate the expression of EMT-inducing factors Escargot and Zfh1 (the homolog of Zeb1), triggering a strong burst of stem cell differentiation in the intestine^48^. Similarly, we found that *mir-8* expression in RNSCs forced *esg+* cells to differentiate and only rare RNSCs persisted after 8 days of treatment (Fig. 3a). Upon *mir-8* expression, the progeny (H2B::RFP-positive, GFP-negative cells) of RNSCs present in lower tubules also expressed Alkaline Phosphatase 4, a marker of a subset of differentiated principal cells^49^ confirming that the depletion of RNSCs upon *mir-8* overexpression was caused by their premature differentiation (Supplementary Fig. 3b). In contrast, no modification of the progenitors/progeny ratio was observed in *svb*-RNAi conditions when compared to controls, showing that *svb* depletion does not trigger RNSC differentiation (Fig. 3a). Finally, we tested whether svb-depleted RNSCs were lost because they underwent apoptosis. Since apoptotic figures are difficult to observe in the digestive track^50^ including the Malpighian tubules, we assayed consequences of blocking programmed cell death by expressing the viral caspase inhibitor p35 ^51^. Although the expression of p35 had no detectable effect by itself on RNSCs, it significantly rescued the number of RNSCs when *svb* was depleted (Fig. 3c and see below).

**Figure 3:**
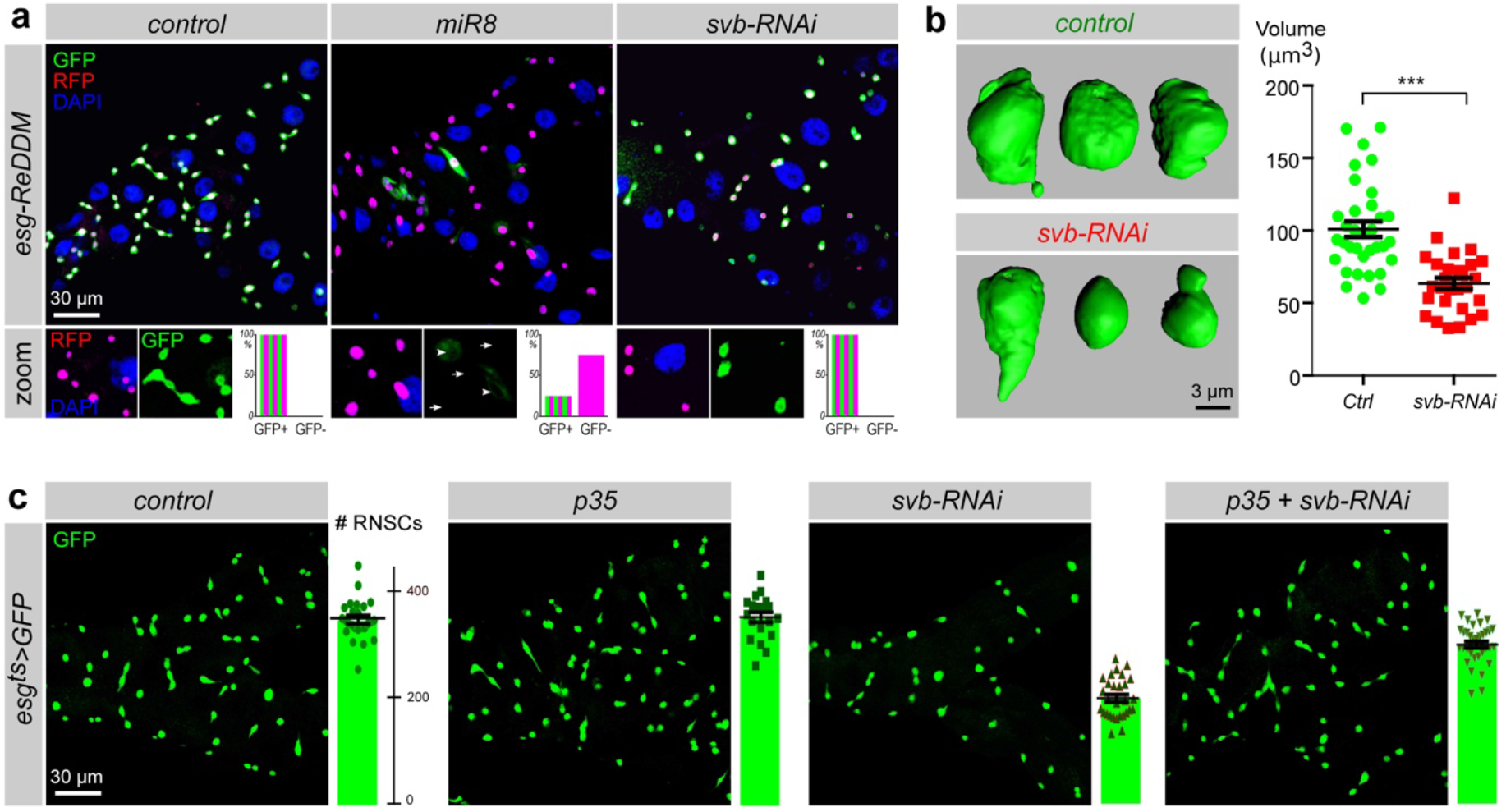
*svb* protects RNSCs from apoptosis. **(a)** Lineage-tracing experiments (*esg*-ReDDM) at 8 days after induction. While RNSCs express both mCD8::GFP (green) and H2B::RFP (red), only the stable H2B::RFP protein persists in their progeny. Nuclei are in blue (DAPI). RFP positive cells that lack or display remnants of GFP levels are indicated by arrows and arrowheads, respectively. **(b)** Left, threedimensional reconstruction of 3 different RNSCs in control (*esg*^*ts*^>*GFP*) and *svb-RNAi* contexts. Pictures show GFP after 8 days of treatment. Right, quantification of the RNSC volume in control (*Ctrl, green*) and *svb-RNAi* conditions. **(c)** Rescue of svb-depleted RNSCs by p35. *esg*^*ts*^ was used to drive the expression of indicated transgenes (together with GFP), during 8 days. Quantification of *esg+* cells is shown at the right of each picture.

Taken together, these data show that the loss of RNSCs observed upon *svb* loss of function is primarily due to stem cell death, indicating that a main role of Svb is to protect adult stem cells from undergoing apoptosis.

### Svb acts downstream of Hippo

During epidermal development, *svb* is expressed in post-mitotic cells where it acts as a terminal differentiation factor that controls cell shape remodeling^15, 52^. We noticed that RNSCs lacking *svb* displayed a reduced size, as well as an altered morphology (Fig. 3b). One could speculate that these defects in cell shape might stress RNSCs and thus induce apoptosis. Indeed, the Hippo pathway^53, 54^ is a key sensor of mechanical stress renowned to induce apoptosis following cytoskeleton modifications^55, 56^. The core Hippo complex is composed of two kinases, Hippo (Hpo) and Warts and two scaffolding proteins, Salvador and Mob As Tumor Suppressor^53, 54^ Activation of Hippo leads to the phosphorylation of the co-transcription factor Yorkie (Yki) that results in Yki nuclear exclusion/degradation, preventing its positive action on the transcription of target genes such as *DIAP1* and *bantam*^53, 54^. Previous work has shown that the Hippo pathway is a key regulator of the *Drosophila* gut homeostasis, controlling survival and proliferation of stem cells for tissue regeneration^57, 58^. Consistently, we found that the activation of Hpo induced a strong reduction in the number of RNSCs. However, co-expression of OvoB, the constitutive activator isoform of Svb, together with Hpo was sufficient to rescue the loss of RNSCs (Fig. 4a). These results therefore demonstrated that if Svb and Hpo interact for the homeostasis of RNSCs, the loss of RNSCs observed upon *svb* knockdown is not due to the activation of the Hippo pathway, since Svb is instead acting downstream Hpo. In contrast, overexpression of Yki (mimicking a loss of Hippo signaling^59^) induced a strong increase in *esg+* renal cells, which displayed abnormal tumor-like morphology when compared to wild-type RNSCs (Fig. 4a). Unexpectedly, this increase in the number of renal stem cells was entirely suppressed upon *svb* depletion, or expression of the constitutive repressor OvoA (Fig. 4a and Supplementary Fig. 3f). Quantification indicated that *esg+* cells overexpressing Yki were even more sensitive to *svb* loss-of-function than otherwise normal RNSCs (Fig. 4a), a result well in line with the extra-resistance of intestinal stem cells to apoptosis when compared to tumorous stem-like cells^50^. Hence, the function of Yki in RNSCs requires Svb, suggesting that Svb was acting either downstream or in parallel with this nuclear effector of Hippo. Several lines of evidence ruled out the former and validated the latter hypothesis. First, knocking down Yki also led to RNSC loss (Supplementary Fig. 3c). Expression of the Svb constitutive activator (OvoB) was nevertheless not able to rescue RNSC survival in the absence of Yki (as opposed to the over expression of Hpo, Fig. 4a), showing that Svb requires Yki activity for RNSC maintenance (Fig. 4a and Supplementary Fig. 3c). Second, although Yki is sufficient to induce *DIAP1* expression^60^ (and see below), Yki was not able to rescue the lack of Svb while DIAP1 alone did (Fig. 4a). Indeed, we found that DIAP1 was sufficient to compensate for svb-depletion (Fig. 4a and Supplementary Fig. 3f). In sum, while both Svb and Yki are required for RNSC maintenance, re-expression of Yki is not sufficient to rescue the loss of *svb* function. Reciprocally, Svb is not sufficient to rescue a proper RNSC compartment in the absence of Yki, showing that Svb and Yki act in parallel for the survival of adult stem cells.

**Figure 4:**
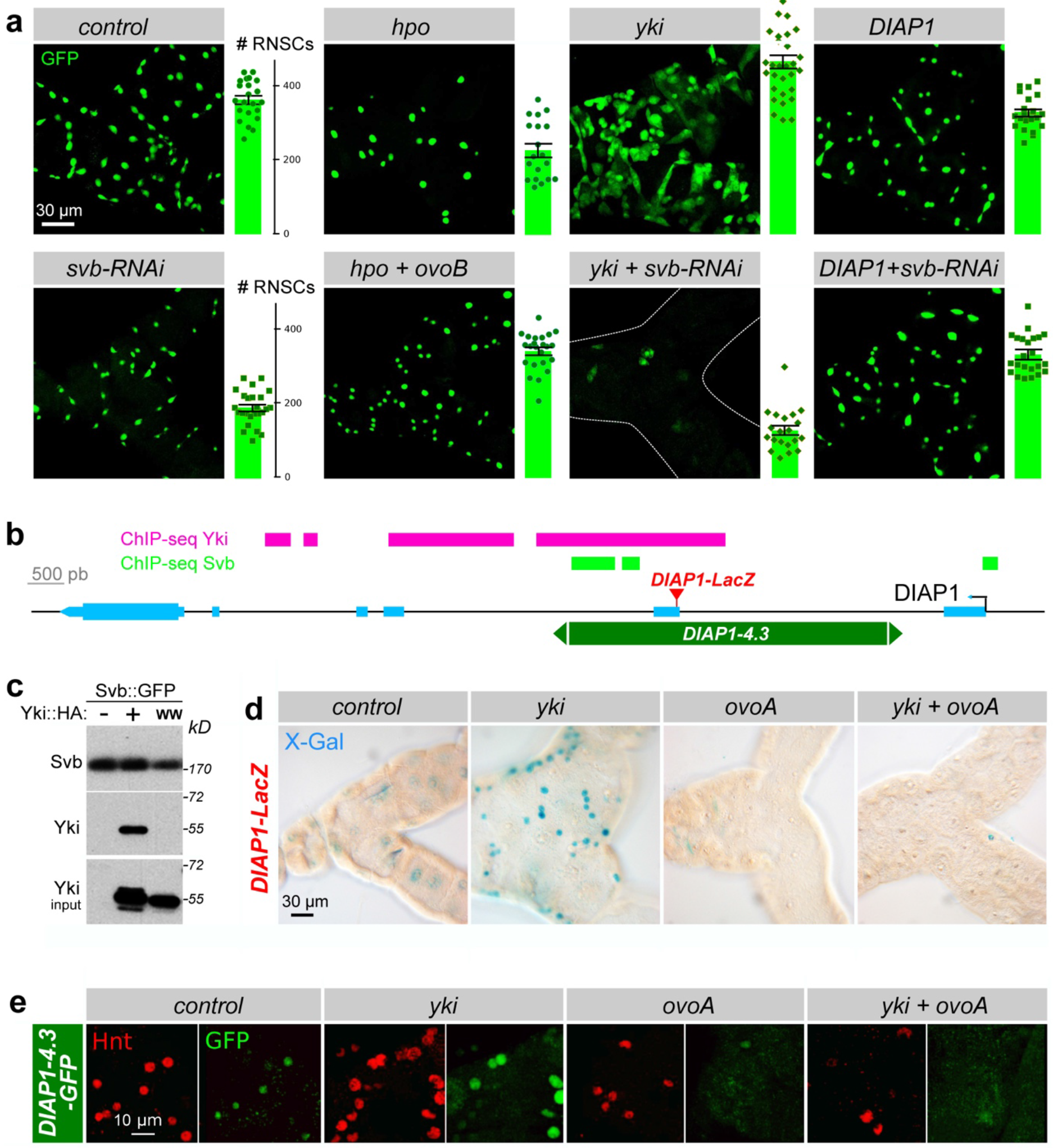
*svb* is a member of the Hippo pathway. **a)** Pictures of Malpighian tubules with *esg*^*ts*^-driven expression of GFP (control) and indicated transgenes. Quantification of esg+ cells is given on each picture (see also Supplementary Figure 3f).**(b)** Drawing of the *DIAP1* locus. Exons are represented in cyan, the *DIAP1-4.3* enhancer in dark green and the insertion site of the *DIAP1-LacZ* reporter (J5C8) is in red. Regions bound in ChIP-seq (MACS peaks) by Svb and Yki are indicated in green and magenta,respectively.**(c)** Svb co-immuno-precipitates with Yki in S2 cells. Svb::GFP and Yki::HA, or a mutated form of Yki substituting the WW domains (YkiWW::HA), were expressed in S2 cells. Protein immuno-precipitated by anti-GFP were blotted with anti-GFP and anti-HA antibodies.**(d)** *esg*^*ts*^ was used to drive the expression of *yki*, *svb*^*REP*^ (*ovoA*), and *yki* together with *svb*^*REP*^ in RNSCs. The expression of *DIAP1* was followed by the activity of *DIAP1-lacZ*. **(e)** Expression of the *DIAP1-4.3-GFP* enhancer was followed by immuno-staining against GFP (green) and Hindsight (Hnt, red) revealing the antagonistic influence of Yki and OvoA on RNSCs.

We thus concluded that Svb acts downstream of Hippo cytoplasmic core components and, together with Yki, both nuclear factors are required to protect RNSCs from apoptosis.

### Svb as a novel nuclear effector of the Hippo pathway

Having established that Svb and Yki genetically interact, we wondered whether the two proteins might interact to control the expression of common target genes, *e.g., DIAP1.* Yki is unable to bind DNA by itself and need to associate to sequence-specific-transcription factors^54^. Interestingly, Yki contains two WW protein domains shown to mediate interaction with partners bearing PPxY motifs (such as Wts^60^, Wbp2^61^ and Mad^59^), and we detected two PPxY motifs within the Svb protein, at position 523 (PPFY) and 881 (PPSY). Co-immunoprecipitation assays showed that Svb bound to the wild type form of Yki, while the mutation of Yki WW motifs was sufficient to abrogate the interaction with Svb (Fig. 4c). A second piece of evidence emerged from the comparison of chromatin immuno-precipitation (ChIP-seq) datasets between Svb^14^ and Yki^62^. We found that Svb and Yki share 836 common genomic binding sites (Supplementary Fig. 4a) and statistical tests established the significance of this overlap (Supplementary Fig. 4b). Interestingly, co-binding of Yki was rare for the direct target genes of Svb identified in the epidermis^12-14^, as illustrated by *shavenoid* or *dusky-like* that both lack Yki binding (Supplementary Fig. 4d,d’). In contrast, Svb was often bound to known Yki target genes, such as *bantam*, *fat*, *piwi* or *nanos*^63^ (Supplementary Fig. 4c,c’). ChIP-seq also revealed that Svb and Yki bound to a same region of *DIAP1*, within an enhancer driving specific expression in adult intestinal stem cells^64^ (Fig. 4b). We therefore tested if Svb might regulate *DIAP1* expression. Although very weak in control conditions, expression of *DIAP1-LacZ* was strongly enhanced upon Yki overexpression. This induction was completely antagonized by OvoA (Fig. 4d). Similar results were obtained with the isolated *DIAP1* enhancer *(DIAP1-4.3-GFP)* containing the binding sites of Yki and Svb, the expression of which was again enhanced by Yki overexpression and abrogated upon counteracting Svb activity (Fig. 4e). These data thus strengthen the conclusion that Svb and Yki functionally interact in RNSCs to prevent apoptosis, at least in part through promoting *DIAP1* expression.

One important question was whether the interaction between Svb and the Hippo pathway also took place in other tissues. The function of Hippo has been initially described in imaginal discs, which give rise to most adult tissues^65^ and Yki overexpression promotes cell proliferation in both wing and eye discs^60^. We tested Svb/Yki genetic interactions in the wing using *collier-Gal4* that drives expression in the medial (L3-L4) intervein region. *Yki* over-expression resulted in the expansion of this region due to tissue overgrowth (Supplementary Fig. 3d). In contrast, OvoA leads to both a reduction of the L3-L4 region and the absence of epidermal trichomes. As in RNSCs, OvoA was epistatic to Yki, since the wing region expressing both *yki* and *ovoA* was smaller than in controls and lacked trichomes (Supplementary Fig. 3d). In the eye, overexpression of Yki using the *GMR-Gal4* driver promoted extra cell proliferation resulting in an increased eye size. Similar results were obtained following *pri* overexpression, and co-expressing *pri* and *yki* resulted in a synergistic eye growth (Supplementary Fig. 3e). Northern blotting of RNAs extracted from adult heads revealed that *DIAP1* mRNA levels were increased following *pri* overexpression (Supplementary Fig. 4e), while there was no effect on *yki* or *cycE* mRNA.

We interpret these results to imply that Svb functionally interacts with Yorkie, both in adult stem cells and in developing tissues, to regulate a subset of transcriptional targets of the Hippo pathway, including the activation of *DIAP1* expression.

## Conclusions

Our results show that Shavenbaby is expressed and required for the maintenance of adult renal stem cells (RNSCs) in flies, supporting the conclusion that the OvoL/Svb family of transcription factors plays a key and evolutionarily-conserved role in the behavior of progenitors-stem cells.

The role of Svb in adult stem cell maintenance in flies requires both a fine control of its expression and of its transcriptional activity. Svb expression in RNSCs involves at least two separable enhancers, driving similar expression patterns. Svb was one the first cases to reveal the functional importance of apparently redundant (or shadow) enhancers in the phenotypic robustness of developmental networks^8^. Our data suggest that a similar *cis*-regulatory architecture is also underlying the control of adult stem cells.

RNSCs maintenance further requires a proper post-translational maturation of the Svb protein, in the response to Pri smORF peptides. During both embryonic^17^ and post-embryonic development^18, 45^, the main role of Pri peptides is to provide a temporal control of Svb activity, conveying systemic steroid signaling^18^. It is therefore possible that Pri smORF peptides also connect genetic networks to hormonal control for the regulation of adult stem cells. Recent work has shown that various smORF peptides contribute to the regulation of developmental pathways, muscle formation and physiology, etc…^16, 66, 67^, and our findings extend their influence to the control of adult stem cells. It has been proposed that the emerging field of smORF peptides may open novel therapeutic strategies, for example peptidomimetic drugs, which might also be of interest for regenerative medicine.

Complementary data establish that a main function of Svb in adult stem cells is mediated by a functional interplay with the Hippo pathway, well established for its roles in the control of adult stem cells^53, 57, 58, 68^. Our results indicate that Svb behaves as a novel nuclear effector of Hippo, relying on a direct interaction with Yorkie in order to protect stem cells from apoptosis, at least in part through the regulation of *DIAP1* expression. Analysis of genome- wide binding events further suggests that the Svb/Yki interaction is involved in the control of a broader set of Hippo-regulated genes, including during development. Since both Pri and Ubr3 are also essential for the survival of adult stem cells, it is interesting to note that Ubr3 protects the DIAP1 protein from degradation^69^, and direct binding of Ubr3 on the activated form of DIAP1 is elicited in the presence of Pri peptides^45^. Therefore, in addition to the control of *DIAP1* expression (*via* Svb), Ubr3 and Pri could also stabilize the DIAP1 protein to protect stem cells from apoptosis. Although initially restricted to TEAD transcription factors, the number of Yorkie (YAP/TAZ) nuclear partner is rapidly growing^54^. Recent work has shown a direct interaction of YAP/TAZ with the Snail/Slug pro-EMT factors in the control of stem cell renewal and differentiation^70^. As previously reported for intestinal stem cells^40, 41, 48^, pro-EMT regulators are also required for preventing premature differentiation of renal stem cells. While pro-EMT and OvoL factors are often viewed as antagonistic factors ^19, 21, 25^, *in vivo* studies in *Drosophila* stem cells show that they both contribute to their maintenance, Svb/Yki preventing their apoptosis and EMT factors their differentiation. Many studies having implicated the Hippo pathway, pro-EMT and OvoL/Svb factors in various tumors, new insights into their functional interactions in adult stem cells may provide additional knowledge directly relevant to understand their connections in human cancers.

## Methods

**Fly stocks.** The following *Drosophila* stocks were used in this study: *tsh-LacZ* (BL#11370), *esg-lacZ* (BL#10359), *esg-Gal4, UAS-GFP; tubulin-Gal80^ts^/SM6-TM6B* ^43^ (B. Edgar), *yw, hsFLP, tubulin-Gal80 FR19A; UAS-mcd8::GFP/Cyo; tubulin-Gal4/TM6 Tb* (N. Tapon*), esg- Gal4,UAS-mcd8::GFP/Cyo; UAS-H2B::RFP, tubulin-Gal80^ts^/TM2 ^48^* (M. Dominguez), *col- Gal4, UAS-mcd8::GFP/Cyo* and *domeMeso-Gal4* (M. Crozatier), *GMR-Gal4 /Cyo* (BL#9146), *tal-Gal4/ TM3 Sb* (J.P. Couso), *svb^E^-GFP, svb^E10^-lacZ, svb^E3N^-lacZ, svb^E6^-lacZ* (D. Stern), *svb^E3N^-GFP* (this work), *svb^R9^, FRT19A/FMO ^44^, ubr3^B^ FRT19A/FMO* ^45^(H. Bellen), *Diap1-lacZ* (BL#12093), *UAS-svb RNAi* (VDRC #v40316), *UAS-ubr3 RNAi* (VRDC #22901), *UAS-yki RNAi* (VDRC #KK104523), *UAS-pri RNAi* (J.P. Couso), *UAS-OvoA* ^44^ *UAS-OvoB* ^44^, *UAS-EcRDN* (BL#9449), *UAS-mir8* (S.M. Cohen), *UAS-p35* (B. Monier), *UAS-hpo/CyO* (N. Tapon), *UAS-yki/TM3 Sb* (D.J. Pan), *UAS-DIAP1* (N. Tapon), *UAS- pri/CyO* (J.P. Couso).

Flies were cultured (unless otherwise noted) at 25°C, using standard cornmeal food (*per* liter: 17g inactivated yeast powder, 80g corn flour, 9g agar, 45g white sugar and 17ml of Moldex). Similar results were also observed using a richer medium (same composition except 45g of yeast powder). Female adult flies were used in all analyses throughout the study and placed on fly food supplemented with fresh yeast, which was changed every two days. Conditional expression in RNSCs was achieved by maintaining *tub-Gal80^ts^* expressing flies at 18°C, until adulthood. Eclosed females aged for 3- to 4-days were shifted to 29°C for induction of gene expression and were kept at 29°C for the indicated period of time (in most cases 8 days). Virgin females bearing *svb^R9^*, *svb^PL107^* or *ubr3^B^* mutations^44, 45^ over FM0 balancers were mated with males of the following genotype: *yw, hs-FLP, tub-Gal80, FRT19A; UAS::mcd8-GFP; tub-Gal4/TM6,Tb.* Females of the correct phenotype (no *B* and no *Tb)* were heat shocked for 1h at 37°C. Flies were transferred on fresh food every two days and dissected at the indicated time. Detailed information about the full genotype of each *Drosophila* stock is given in the genotype section below.

**Histology.** Tissues were dissected in 1X PBS, fixed in 4% formaldehyde in PBS for 15 min at room temperature, washed for 5 min in PBS containing 0.1% Triton X-100 (PBT) and fixed again during 20 min. Following a 5 min wash (PBT 0.1), tissues were then blocked for 30 minutes in PBT containing 1% BSA. Primary antibodies were incubated overnight at 4°C. Anti-ß-Galactosidase (Cappel) antibody was used at 1/1000, Cut (DSHB) and GFP antibodies at 1/200, and phospho-HistoneH3 (Upstate) and Disc-large (DSHB) antibodies at 1/500. AlexaFluor-488 or 555 secondary antibodies (Molecular Probes) were incubated for 2 hrs at room temperature at 1/500. After washes, tissues were mounted in Vectashield (Vector). For X-gal staining, adult tissues were dissected in 1X PBS, fixed in 1% glutaraldehyde in PBS for 15 min at room temperature and washed in PBS. The staining solution was warmed up at 37°C for 10 min plus 10 other min after addition of 8% X-Gal (5-bromo-4-chloro-3-indoyl-ß- D-Galactopyranoside). The X-Gal solution used to reveal the ß-Galactosidase activity was: 10mM NaH2PO4.H2O/ NA2HPO4.2H2O (pH=7.2), 150mM NaCl, 1mM MgCl2.6H2O, 3.1mM K4 (FeII(CN)6), 3.1mM K3 (FeIII(CN)6), 0.3 % Triton X-100. Bright-field pictures were acquired using a Nikon eclipse 90i microscope.

**Microscopy, image and statistical analysis.** Images of whole Malpighian tubules were acquired on a LSM710 confocal scanning microscope (X20 objective), using automated multi-position scan. After stitching, tiled images of individual pairs of tubules were analyzed with IMARIS 8.0 to quantify the number of GFP-positive cells. Data of at least three independent experiments (approx. 20 tubules) were analyzed with GraphPad Prims 5, using two-tailed Mann-Whitney tests. The statistical significance of differences observed between compared genotypes was indicated as: *** (p<0.001), ns (p>0.05). Close-up pictures were acquired using Leica SPE and Leica SP8 confocal laser scanning microscopes (X40 and X63 objectives). Laser intensity and background filtering was set according to the control samples and remained the same for all subsequent samples. The intensity of most pictures has been enhanced equally for all images within the same experiment using adjustments in Photoshop CS5. All images were processed using Adobe Photoshop, Illustrator CS5 and Inkscape 0.91.

**Western blotting and immunoprecipitation**. *Drosophila* S2 cells were grown in Schneider medium supplemented with 10% fetal calf serum and 1% penicillin/streptomycin (Invitrogen) at 25°C. We used stable cell lines co-expressing the copper-inducible constructs pMT-Svb::GFP and pMT-pri^17^. S2 cell lines were cultured in six-well plates (1.75x106 cells/3ml) and transfected in 100 μl of Opti-MEM (Invitrogen) with 3 μl of FugeneHD (Promega) and the indicated constructs. CuSO4 (final concentration of 1mM) was used to induce the expression of pMT plasmids. The following plasmids were used: pAc-Yki::HA and its related mutated version pAc-Yki-WW::HA. Cells were lysed in 250 μl of ice-cold lysis buffer (150 mM NaCl, 50mM Tris [pH 8], 0.5% NP40, 1mM EGTA, 0.5M NaF, 200mM vanadate, phosphatase inhibitor cocktail 1 (Sigma) and protease inhibitors (Roche). After clearing by centrifugation at 14,000 rpm for 10 min, immuno-precipitations were done from transfected lysates in lysis buffer using anti-GFP antibody (GFP-trap, Chromotek). Immuno-precipitated samples were separated by SDS-PAGE and transferred to PVDF membranes, then blotted using anti-GFP (TP401, Acris Antibodies, 1:10000) and anti-HA (Covance, 1:2.000) antibodies. Secondary antibodies anti-mouse or anti-rabbit IgG-HRP conjugates (Jackson Laboratory, 1:10.000) were detected using ECL Clarity (Biorad).

**Northern blot analysis.** Using adult total RNAs as a starting material, DNA fragments containing coding sequence of *yki, CycE* and *DIAP1* were reverse transcribed and PCR-amplified with pairs of specific primers: CTGCC CAACT CCTTC TTCAC (forward) and AACTG AATGG GGCTG ATGAC (reverse) for *yki*; GATGA CGTTG AGGAG GAGGA (forward) and TGCGT CTTCT GCACC TTATG (reverse) for *CycE*; CCGAG GAACC TGAAA CAGAA (forward) and GCACA ACTTT TCCTC GGGTA (reverse) for *DIAP1.* A SP6 promoter sequence (CAAGC TATTT AGGTG ACACT ATAG) was attached to each reverse primer for *in vitro* transcription. DIG-labelled probes were prepared with SP6 RNA polymerase, according to the supplier’s manual (Roche). Northern blot analysis was described previously^18^. Briefly, 2 days-old adults were frozen with liquid nitrogen and heads were sorted with sieves, followed by RNA purification with Isogen (Nippon Gene). 1 μg of RNA *per* lane was separated by formaldehyde-agarose gel electrophoresis and then transferred to a nylon membrane (Roche). Hybridization and wash procedures were carried out at 52°C and 65°C, respectively. The filters were reacted with an alkaline phosphatase-conjugated anti-DIG antibody (Roche) and chemiluminescent reactions with CPD-Star (Roche) were detected by LAS 4000mini (GE Healthcare).

## Genotypes

**Figure 1A:** *yw/w; esg-Gal4, UAS-GFP/ tsh-LacZ*

**Figure 1B:** *yw/w; esg-LacZ/+; svb^E3N^-GFP*/+

**Figure 1C&D *control:*** *yw/w; esg-Gal4, UAS-GFP*/+; *tubulinGal80^ts^*/+

**Figure 1C&D *svb-RNAi:*** *yw/w; esg-Gal4, UAS-GFP*/+*; tubulin-Gal80^ts^*/ *UAS-svb-RNAi*

**Figure 1E *control:*** *yw, hsFLP, tubulin-Gal80 FR19A; UAS-mCD8::GFP*/+*; tubulin-Gal4/ry^506^*

**Figure 1E *svb*^*R9*^:** *yw, hsFLP, tubulin-Gal80 FR19A/yw svb^R9^ FRT19A; UAS-mCD8::GFP*/+*; tubulin-Gal4*/+

**Figure 2B:** *tal-Gal4/ UAS-HB2::RFP*

**Figure 2C *control*:** *yw, hsFLP, tubulin-Gal80 FR19A; UAS-mCD8::GFP*/+*; tubulin-Gal4/ry^506^*

**Figure 2C *ubr3^B^:*** *yw, hsFLP, tubulin-Gal80 FR19A/yw ubr3^B^ FRT19A; UAS-mCD8::GFP*/+*; tubulin-Gal4*/+

**Figure 2D *control*:** *yw/w; esg-Gal4, UAS-GFP*/+*; tubulin-Gal80^ts^*/+

**Figure 2D *svb-RNAi*:** *yw/w; esg-Gal4, UAS-GFP*/+*; tubulin-Gal80^ts^/ UAS-svb-RNAi*

**Figure 2D *ovoA*:** *yw/w; esg-Gal4, UAS-GFP*/+*; tubulin-Gal80^ts^/ UAS-ovoA*

**Figure 2D *ovoB*:** *yw/w; esg-Gal4, UAS-GFP*/+?*; tubulin-Gal80^ts^/ UAS-ovoB*

**Figure 2D *pri-RNAi:*** *yw/w; esg-Gal4, UAS-GFP/+; tubulin-Gal80^ts^/ UAS-pri-RNAi*

**Figure 2D *EcRDN*:** *yw/w; esg-Gal4, UAS-GFP/+; tubulin-Gal80^*ts*^/ UAS-EcRDN^B2w650A^*

**Figure 2D *ubr3-RNAi*:** *yw/w; esg-Gal4, UAS-GFP*/ *UAS-ubr3 RNAi; tubulin-Gal80^ts^*/+

**Figure 2D *ubr3-RNAi* + *ovoB*:** *yw/w; esg-Gal4, UAS-GFP/ubr3 RNAi; tubulin-Gal80^ts^*/ *UAS-ovoB*/+

**Figure 3A *control:*** *yw/w; esg-Gal4, UAS-mCD8::GFP/+; UAS-H2B::RFP, tubulin-Gal80^ts^/+*

**Figure 3A *mir8:*** *yw/w, UAS-mir8; esg-Gal4, UAS-mCD8::GFP/+; UAS-H2B::RFP, tubulin- Gal80^ts^/+*

**Figure 3A *svb-RNAi:*** *yw/w; esg-Gal4, UAS-mCD8::GFP/+; UAS-H2B::RFP, tubulin- Gal80^ts^/ UAS-svb RNAi*

**Figure 3B *control:*** *yw/w; esg-Gal4, UAS-GFP/+; tubulin-Gal80^ts^/+*

**Figure 3B *svb-RNAi:*** *yw/w; esg-Gal4, UAS-GFP/+; tubulin-Gal80^ts^/ UAS-svb-RNAi*

**Figure 3C *control:*** *yw/; esg-Gal4, UAS-GFP/+; tubulin-Gal80^ts^/+*

**Figure 3C *p35:*** *yw/; esg-Gal4, UAS-GFP/ UAS-p35; tubulin-Gal80^ts^/+*

**Figure 3C *svb-RNAi:*** *yw/; esg-Gal4, UAS-GFP/+; tubulin-Gal80^ts^/ UAS-svb-RNAi*

**Figure 3C *p35+ svb-RNAi:*** *yw/; esg-Gal4, UAS-GFP/ UAS-p35; tubulin-Gal80^ts^/ UAS-svb-RNAi*

**Figure 4A *control:*** *yw/; esg-Gal4, UAS-GFP/+; tubulin-Gal80^ts^/+*

**Figure 4A *hpo:*** *yw/; esg-Gal4, UAS-GFP/ UAS-hpo; tubulin-Gal80^ts^/+*

**Figure 4A *yki*:** *yw/; esg-Gal4, UAS-GFP/+; tubulin-Gal80^ts^/ UAS-yki*

**Figure 4A *DIAP1:*** *yw/; esg-Gal4, UAS-GFP/ +; tubulin-Gal80^ts^/ UAS-DIAP1*

**Figure 4A *svb-RNAi:*** *yw/; esg-Gal4, UAS-GFP/ +; tubulin-Gal80^ts^/ UAS-svb-RNAi*

**Figure 4A *hpo+ ovoBt:*** *yw/; esg-Gal4, UAS-GFP/ UAS-hpo; tubulin-Gal80^ts^/ UAS-ovoB*

**Figure 4A *yki+ svb-RNAi:*** *yw/; esg-Gal4, UAS-GFP/ +; tubulin-Gal80^ts^/ UAS-ovoB, UAS-yki*

**Figure 4A *DIAP1+ svb-RNAi:*** *yw/; esg-Gal4, UAS-GFP/ +; tubulin-Gal80^ts^/ UAS-DIAP1, UAS-svb-RNAi*

**Figure 4D *control:*** *yw/; esg-Gal4, UAS-GFP/ +; tubulin-Gal80^ts^/ DIAP1-LacZ/+*

**Figure 4D *yki:*** *yw/; esg-Gal4, UAS-GFP/ +; tubulin-Gal80^ts^/ DIAP1-LacZ/ UAS-yki*

**Figure 4D ovoA:** *yw/; esg-Gal4, UAS-GFP/ +; tubulin-Gal80^ts^/DIAP1-LacZ/ UAS-ovoA*

**Figure 4D *yki + ovoA:*** *yw/; esg-Gal4, UAS-GFP/ +; tubulin-Gal80^ts^/ DIAP1-LacZ/ UAS-yki, UAS-ovoA*

**Figure 4E *control:*** *yw/; esg-Gal4, UAS-GFP/ +; tubulin-Gal80^ts^/DIAP14.3-GFP/+*

**Figure 4E *yki:*** *yw/; esg-Gal4, UAS-GFP/ +; tubulin-Gal80^ts^/ DIAP14.3-GFP/ UAS-yki*

**Figure 4E ovoA:** *yw/; esg-Gal4, UAS-GFP/ +; tubulin-Gal80^ts^/DIAP14.3-GFP/ UAS-ovoA*

**Figure 4E *yki + ovoA:*** *yw/; esg-Gal4, UAS-GFP/ +; tubulin-Gal80^ts^/ DIAP14.3-GFP/ UAS- yki, UAS-ovoA*

**Figure Sup lB:** *yw/w; esg-LacZ/svb^E10^-GFP*

**Figure Sup lC:** *yw/w; esg-LacZ/+*

**Figure Sup lC:** *w; svb^E3N^-LacZ/+*

**Figure Sup lC:** *w; svb^E6^-LacZ/+*

**Figure Sup lD:** *yw/w; esg-Gal4, UAS-GFP/+*

**Figure Sup lE *control:*** *yw/w; dome-meso-Gal4, UAS-mCherry/+*

**Figure Sup lE *svb-RNAi:*** *yw/w; dome-meso-Gal4, UAS-mCherry/+; UAS-svb-RNAi/+*

**Figure Sup 2B:** *yw/w; esg-LacZ/+*

**Figure Sup 2B:** *w; priA-LacZ/+*

**Figure Sup 2B**: *yw/+; priJ-LacZ/+*

**Figure Sup 3A:** *yw/w; esg-Gal4, UAS-GFP/+*

**Figure Sup 3B control:** *yw/w; esg-Gal4, UAS-mCD8::GFP/+; UAS-H2B::RFP, tubulin- Gal80^ts^/+*

**Figure Sup3B miR-8:** *yw/UAS-mir8; esg-Gal4, UAS-mCD8::GFP/+; UAS-H2B::RFP, tubulin-Gal80^ts^/+*

**Figure Sup3C *control:*** *yw/; esg-Gal4, UAS-GFP/+; tubulin-Gal80^ts^/ +*

**Figure Sup3C *ovoB:*** *yw/; esg-Gal4, UAS-GFP/+; tubulin-Gal80^ts^/UAS-ovoB*

**Figure Sup3C *yki-RNAi:*** *yw/; esg-Gal4, UAS-GFP/+; tubulin-Gal80^ts^/ UAS-yki*

**Figure Sup3C *yki-RNAi + ovoB:*** *yw/; esg-Gal4, UAS-GFP/+; tubulin-Gal80^ts^/ UAS-yki, UAS-ovoB*

**Figure Sup3D *ctrl:*** *yw/w; col-Gal4, UAS-mCD8::GFP/+*

**Figure Sup3D *yki-RNAi:*** *yw/w; col-Gal4, UAS-mCD8::GFP/+; UAS-yki/+*

**Figure Sup3D ovoA:** *yw/w; col-Gal4, UAS-mCD8::GFP/+; UAS-ovoA/+*

**Figure Sup3D *yki-RNAi* + *ovoA*:** *yw/w; col-Gal4, UAS-mCD8::GFP/+; UAS-yki, UAS- ovoA/+*

**Figure Sup3E *ctrl:*** *yw/w; GMR-Gal4/+*

**Figure Sup3E *pri:*** *yw/w; GMR-Gal4/UAS-pri*

**Figure Sup3E yki:** *yw/w; GMR-Gal4/+; UAS-yki*

**Figure Sup3E *pri+ yki:*** *yw/w; GMR-Gal4/UAS-pri; UAS-yki/+*

## Acknowledgments

We are grateful to FlyBase, Bloomington and Vienna stock centers, Developmental Studies Hybridoma Bank, as well as N. Tapon, J. Colombani, K.F. Harvey, D.J. Pan, J. Dow and H. Skaer and M. Crozatier, for providing flies, antibodies and molecular reagents. We also thank
B. Ronsin (Toulouse RIO Imaging) for help with microscopy, A. Alsawadi, A. Dib, J. Zanet, M. Soulard and P. Valenti for experimental support. We also thank all members of the Payre lab for critical reading of the manuscript. This work was supported by ANR (Chrononet), Fondation pour le Recherche Médicale (FRM), Programme Scientifique de Cooperation Internationale (PSCI) for D O. and F.P., and by MEXT KAKENHI (JP26113006) for Y.K.. J.B. was supported by fellowships from “Ministére de l’Enseignement et de la Recherche” and “La Ligue contre le Cancer.”

## Author Contributions

C. P., S.P., Y.K., D.O. and F.P. conceived and directed the project, following initial observations made by D.O.. J.B. carried out most of the experiments presented here, under the supervision of C.P. and other experiments were conducted by C.P., K.A., Y.Y, S.I and D.O.. A.M.F analyzed NGS data. J.B., C.P., K.A., Y.Y., S.I., H.C.D., S.P., Y.K., D.O. and F.P. analyzed data and contributed to their interpretation. J.B., C.P. and F.P. prepared the figures and wrote the manuscript. All authors helped to write the paper.

## References

1. Kumar, A. et al. Molecular phylogeny of OVOL genes illustrates a conserved C2H2 zinc finger domain coupled by hypervariable unstructured regions. PLoS One 7, e39399 (2012).

2. Mevel-Ninio, M., Terracol, R., Salles, C., Vincent, A. & Payre, F. ovo, a Drosophila gene required for ovarian development, is specifically expressed in the germline and shares most of its coding sequences with shavenbaby, a gene involved in embryo patterning. Mech Dev 49, 83–95 (1995).

3. Payre, F., Vincent, A. & Carreno, S. ovo/svb integrates Wingless and DER pathways to control epidermis differentiation. Nature 400, 271–275 (1999).

4. Dai, X. et al. The ovo gene required for cuticle formation and oogenesis in flies is involved in hair formation and spermatogenesis in mice. Genes Dev 12, 3452–3463 (1998).

5. Li, B. et al. Ovol1 regulates meiotic pachytene progression during spermatogenesis by repressing Id2 expression. Development 132, 1463–1473 (2005).

6. Wells, J. et al. Ovol2 suppresses cell cycling and terminal differentiation of keratinocytes by directly repressing c-Myc and Notch1. J Biol Chem 284, 29125–29135 (2009).

7. Crocker, J. et al. Low affinity binding site clusters confer hox specificity and regulatory robustness. Cell 160, 191–203 (2015).

8. Frankel, N. et al. Phenotypic robustness conferred by apparently redundant transcriptional enhancers. Nature 466, 490–493 (2010).

9. Frankel, N. et al. Morphological evolution caused by many subtle-effect substitutions in regulatory DNA. Nature 474, 598–603 (2011).

10. McGregor, A.P. et al. Morphological evolution through multiple cis-regulatory mutations at a single gene. Nature 448, 587–590 (2007).

11. Sucena, E., Delon, I., Jones, I., Payre, F. & Stern, D.L. Regulatory evolution of shavenbaby/ovo underlies multiple cases of morphological parallelism. Nature 424, 935–938 (2003).

12. Chanut-Delalande, H., Fernandes, I., Roch, F., Payre, F. & Plaza, S. Shavenbaby couples patterning to epidermal cell shape control. PLoS Biol 4, e290 (2006).

13. Fernandes, I. et al. Zona pellucida domain proteins remodel the apical compartment for localized cell shape changes. Dev Cell 18, 64–76 (2010).

14. Menoret, D. et al. Genome-wide analyses of Shavenbaby target genes reveals distinct features of enhancer organization. Genome Biol 14, R86 (2013).

15. Chanut-Delalande, H., Ferrer, P., Payre, F. & Plaza, S. Effectors of tridimensional cell morphogenesis and their evolution. Semin Cell Dev Biol 23, 341–349 (2012).

16. Zanet, J., Chanut-Delalande, H., Plaza, S. & Payre, F. Small Peptides as Newcomers in the Control of Drosophila Development. Curr Top Dev Biol 117, 199–219 (2016).

17. Kondo, T. et al. Small peptides switch the transcriptional activity of Shavenbaby during Drosophila embryogenesis. Science 329, 336–339 (2010).

18. Chanut-Delalande, H. et al. Pri peptides are mediators of ecdysone for the temporal control of development. Nat Cell Biol 16, 1035–1044 (2014).

19. Watanabe, K. et al. Mammary morphogenesis and regeneration require the inhibition of EMT at terminal end buds by Ovol2 transcriptional repressor. Dev Cell 29, 59–74 (2014).

20. Roca, H. et al. A bioinformatics approach reveals novel interactions of the OVOL transcription factors in the regulation of epithelial - mesenchymal cell reprogramming and cancer progression. BMC Syst Biol 8, 29 (2014).

21. Hong, T. et al. An Ovol2-Zeb1 Mutual Inhibitory Circuit Governs Bidirectional and Multi-step Transition between Epithelial and Mesenchymal States. PLoS Comput Biol 11, e1004569 (2015).

22. Roca, H. et al. Transcription factors OVOL1 and OVOL2 induce the mesenchymal to epithelial transition in human cancer. PLoS One 8, e76773 (2013).

23. Wang, Z.H. et al. Ovol2 gene inhibits the Epithelial-to-Mesenchymal Transition in lung adenocarcinoma by transcriptionally repressing Twist1. Gene 600, 1–8 (2017).

24. Ricketts, C.J. et al. Genome-wide CpG island methylation analysis implicates novel genes in the pathogenesis of renal cell carcinoma. Epigenetics 7, 278–290 (2012).

25. Jia, D. et al. OVOL guides the epithelial-hybrid-mesenchymal transition. Oncotarget 6, 15436–15448 (2015).

26. Li, S. & Yang, J. Ovol proteins: guardians against EMT during epithelial differentiation. Dev Cell 29, 1–2 (2014).

27. Nieto, M.A., Huang, R.Y., Jackson, R.A. & Thiery, J.P. Emt: 2016. Cell 166, 21–45 (2016).

28. Jolly, M.K. et al. Coupling the modules of EMT and stemness: A tunable ‘stemness window’ model. Oncotarget 6, 25161–25174 (2015).

29. Wu, R.S. et al. OVOL2 antagonizes TGF-beta signaling to regulate epithelial to mesenchymal transition during mammary tumor metastasis. Oncotarget 8, 39401–39416 (2017).

30. Lapan, S.W. & Reddien, P.W. Transcriptome analysis of the planarian eye identifies ovo as a specific regulator of eye regeneration. Cell Rep 2, 294–307 (2012).

31. Lee, B. et al. Transcriptional mechanisms link epithelial plasticity to adhesion and differentiation of epidermal progenitor cells. Dev Cell 29, 47–58 (2014).

32. Kitazawa, K. et al. OVOL2 Maintains the Transcriptional Program of Human Corneal Epithelium by Suppressing Epithelial-to-Mesenchymal Transition. Cell Rep 15, 1359–1368 (2016).

33. Kim, J.Y. et al. OVOL2 is a critical regulator of ER71/ETV2 in generating FLK1+, hematopoietic, and endothelial cells from embryonic stem cells. Blood 124, 2948–2952 (2014).

34. Beyenbach, K.W., Skaer, H. & Dow, J.A. The developmental, molecular, and transport biology of Malpighian tubules. Annu Rev Entomol 55, 351–374 (2010).

35. Denholm, B. Shaping up for action: the path to physiological maturation in the renal tubules of Drosophila. Organogenesis 9, 40–54 (2013).

36. Sözen, M.A., Armstrong, J.D., Yang, M., Kaiser, K. & Dow, J.A. Functional domains are specified to single-cell resolution in a Drosophila epithelium. Proc. Natl. Acad. Sci. U.S.A. 94, 5207–5212 (1997).

37. Singh, S.R., Liu, W. & Hou, S.X. The adult Drosophila malpighian tubules are maintained by multipotent stem cells. Cell Stem Cell 1, 191–203 (2007).

38. Takashima, S., Paul, M., Aghajanian, P., Younossi-Hartenstein, A. & Hartenstein, V. Migration of Drosophila intestinal stem cells across organ boundaries. Development 140, 1903–1911 (2013).

39. Micchelli, C.A. & Perrimon, N. Evidence that stem cells reside in the adult Drosophila midgut epithelium. Nature 439, 475–479 (2006).

40. Loza-Coll, M.A., Southall, T.D., Sandall, S.L., Brand, A.H. & Jones, D.L. Regulation of Drosophila intestinal stem cell maintenance and differentiation by the transcription factor Escargot. EMBO J 33, 2983–2996 (2014).

41. Korzelius, J. et al. Escargot maintains stemness and suppresses differentiation in Drosophila intestinal stem cells. EMBO J 33, 2967–2982 (2014).

42. Stern, D.L. & Frankel, N. The structure and evolution of cis-regulatory regions: the shavenbaby story. Philos. Trans. R. Soc. Lond., B, Biol. Sci. 368, 20130028 (2013).

43. Jiang, H. et al. Cytokine/Jak/Stat Signaling Mediates Regeneration and Homeostasis in the Drosophila Midgut. Cell 137, 1343–1355 (2009).

44. Delon, I., Chanut-Delalande, H. & Payre, F. The Ovo/Shavenbaby transcription factor specifies actin remodelling during epidermal differentiation in Drosophila. Mech Dev 120, 747–758 (2003).

45. Zanet, J. et al. Pri sORF peptides induce selective proteasome-mediated protein processing. Science 349, 1356–1358 (2015).

46. Galindo, M.I., Pueyo, J.I., Fouix, S., Bishop, S.A. & Couso, J.P. Peptides encoded by short ORFs control development and define a new eukaryotic gene family. PLoS Biol 5, e106 (2007).

47. Andrews, J. et al. OVO transcription factors function antagonistically in the Drosophila female germline. Development 127, 881–892 (2000).

48. Antonello, Z.A., Reiff, T., Esther, B.-I. & Dominguez, M. Robust intestinal homeostasis relies on cellular plasticity in enteroblasts mediated by miR-8-Escargot switch. EMBO J. 34, 2025–2041 (2015).

49. Yang, M.Y., Wang, Z., MacPherson, M., Dow, J.A. & Kaiser, K. A novel Drosophila alkaline phosphatase specific to the ellipsoid body of the adult brain and the lower Malpighian (renal) tubule. Genetics 154, 285–297 (2000).

50. Ma, M. et al. Wildtype adult stem cells, unlike tumor cells, are resistant to cellular damages in Drosophila. Dev Biol 411, 207–216 (2016).

51. Hay, B.A., Wolff, T. & Rubin, G.M. Expression of baculovirus P35 prevents cell death in Drosophila. Development 120, 2121–2129 (1994).

52. Payre, F. Genetic control of epidermis differentiation in Drosophila. Int J Dev Biol 48, 207–215 (2004).

53. Pan, D. The hippo signaling pathway in development and cancer. Dev. Cell 19, 491–505 (2010).

54. Staley, B.K. & Irvine, K.D. Hippo signaling in Drosophila: recent advances and insights. Dev. Dyn. 241, 3–15 (2012).

55. Sansores-Garcia, L. et al. Modulating F-actin organization induces organ growth by affecting the Hippo pathway. EMBO J 30, 2325–2335 (2011).

56. Gaspar, P., Holder, M.V., Aerne, B.L., Janody, F. & Tapon, N. Zyxin antagonizes the FERM protein expanded to couple f-actin and yorkie-dependent organ growth. Current Biology 25, 679–689 (2015).

57. Shaw, R.L. et al. The Hippo pathway regulates intestinal stem cell proliferation during Drosophila adult midgut regeneration. Development 137, 4147–4158 (2010).

58. Staley, B.K. & Irvine, K.D. Warts and yorkie mediate intestinal regeneration by influencing stem cell proliferation. Current Biology 20, 1580–1587 (2010).

59. Oh, H. & Irvine, K.D. Cooperative regulation of growth by Yorkie and Mad through bantam. Dev Cell 20, 109–122 (2011).

60. Huang, J., Wu, S., Barrera, J., Matthews, K. & Pan, D. The Hippo signaling pathway coordinately regulates cell proliferation and apoptosis by inactivating Yorkie, the Drosophila homolog of YAP. Cell 122, 421 –434 (2005).

61. Zhang, X., Milton, C.C., Poon, C.L., Hong, W. & Harvey, K.F. Wbp2 cooperates with Yorkie to drive tissue growth downstream of the Salvador-Warts-Hippo pathway. Cell Death Differ. 18, 1346–1355 (2011).

62. Oh, H. et al. Genome-wide association of Yorkie with chromatin and chromatin-remodeling complexes. Cell Rep 3, 309–318 (2013).

63. Zhang, C. et al. The ecdysone receptor coactivator Taiman links Yorkie to transcriptional control of germline stem cell factors in somatic tissue. Dev Cell 34, 168–180 (2015).

64. Poernbacher, I., Baumgartner, R., Marada, S.K., Edwards, K. & Stocker, H. Drosophila Pez acts in Hippo signaling to restrict intestinal stem cell proliferation. Curr Biol 22, 389–396 (2012).

65. Tapon, N. et al. salvador Promotes both cell cycle exit and apoptosis in Drosophila and is mutated in human cancer cell lines. Cell 110, 467–478 (2002).

66. Payre, F. & Desplan, C. RNA. Small peptides control heart activity. Science 351, 226–227 (2016).

67. Pueyo, J.I., Magny, E.G. & Couso, J.P. New Peptides Under the s(ORF)ace of the Genome. Trends Biochem Sci 41, 665–678 (2016).

68. Karpowicz, P., Perez, J. & Perrimon, N. The Hippo tumor suppressor pathway regulates intestinal stem cell regeneration. Development 137, 4135–4145 (2010).

69. Huang, Q. et al. Ubr3 E3 ligase regulates apoptosis by controlling the activity of DIAP1 in Drosophila. Cell Death Differ 21, 1961–1970 (2014).

70. Tang, Y., Feinberg, T., Keller, E.T., Li, X.Y. & Weiss, S.J. Snail/Slug binding interactions with YAP/TAZ control skeletal stem cell self-renewal and differentiation. Nat Cell Biol 18, 917–929 (2016).

